# PfAMA1-expressing chimeric rodent malaria parasites provide an in vivo platform for evaluating multistage interventions against malaria

**DOI:** 10.64898/2026.08.28.747760

**Authors:** Quratul-ain Issahaque, Naoaki Shinzawa, Yuto Kegawa, Takashi Sekine, Hisako Amino, Motomi Torii, Moriya Tsuji, Tomoko Ishino

**Author notes:** Corresponding authors: (NS) (TI).

## Abstract

Apical membrane antigen 1 (AMA1) is expressed in the merozoite and sporozoite infectious stages of the malaria parasite, and upon secretion plays essential roles during host cell invasion. AMA1 is a leading candidate for vaccine development, although specific antibodies frequently fail to inhibit the growth of field-isolated *Plasmodium falciparum* malaria parasites, likely due to the high diversity of polymorphisms in AMA1 surface antigens. A key step for efficient invasion of target cells is tight junction formation through interaction of merozoite-surface AMA1 and rhoptry neck protein 2 (RON2), which is secreted and embedded within the erythrocyte membrane. Antibodies or reagents that disrupt the AMA1-RON2 interaction represent interventions to reduce parasite transmission to humans, as well as to repress clinical symptoms. To create a mouse model system for the evaluation of reagents against *P. falciparum* AMA1 (PfAMA1), we generated CRISPR/Cas9-engineered rodent malaria parasites in which the endogenous *Plasmodium berghei* AMA1 (PbAMA1) was replaced with PfAMA1, resulting in a chimeric line Pb_PfAMA1. Pb_PfAMA1 parasites infect mouse liver and erythrocytes as efficiently as the parental line, demonstrating that PfAMA1 functionally complements the essential roles of PbAMA1. AlphaFold-based structure modeling suggested structural compatibility of the heterologous PfAMA1-PbRON2 interaction, and co-immunoprecipitation analyses supported the functional association of the PfAMA1 and PbRON complex required for merozoite invasion of erythrocytes. Utilizing the interaction-inhibitor R1 peptide with Pb_PfAMA1 sporozoites, we demonstrated that the AMA1-RON2 interaction is crucial for sporozoite invasion of hepatocytes. Repeated infection with Pb_PfAMA1 elicited PfAMA1-reactive antibodies, and immune sera inhibited the growth of the *P. falciparum* lines Pf3D7 and PfHB3B; suggesting that naturally processed parasite-derived PfAMA1 induces antibodies which recognize conserved conformational epitopes. To expand this platform, we replaced circumsporozoite protein PbCSP with PfCSP, to generate dual-chimeric Pb_PfCSP+PfAMA1 parasites. Together, these chimeric parasites establish an in vivo platform for evaluating multistage and multi-antigen interventions against malaria.

**Author summary:** Malaria remains a major global health problem despite the recent introduction of the first WHO-recommended malaria vaccine. The vaccine targets sporozoites before they establish liver infection, and interventions are needed which act at additional stages of the parasite life cycle. Apical membrane antigen 1 (AMA1) is a promising candidate because it is expressed in both merozoites and sporozoites, and participates in host-cell infection at both stages. To enable in vivo evaluation of PfAMA1-targeted interventions, we generated genetically engineered rodent malaria parasites expressing *Plasmodium falciparum* AMA1 in place of the endogenous protein. The chimeric parasites infected mouse livers and erythrocytes comparable to the parental line, demonstrating functional complementation by PfAMA1 in the rodent malaria parasite. Blocking the AMA1–RON2 interaction inhibited sporozoite infection of the liver, highlighting the importance of this interaction at multiple stages of infection. Repeated infection with the chimeric parasite induced antibodies that reduced the growth of two genetically distinct *P. falciparum* strains in vitro, suggesting recognition of native conformational epitopes. This chimeric parasite model provides a practical in vivo platform for developing and evaluating multistage malaria interventions targeting AMA1 and its interaction with RON2.

## Introduction

Malaria continues to be a devastating disease of public health concern, causing more than 600 thousand deaths annually [1]. The disease is caused by *Plasmodium* species, with *Plasmodium falciparum* (*Pf*) being the most virulent. In 2021, WHO recommended the use of the first infection-blocking vaccine, RTS,S, targeting the sporozoite circumsporozoite protein, for African children [1]. Recently, it was reported that among children that are age-eligible for vaccinations the vaccine reduced the severity and lethality by only 30% and 13%, respectively [2,3]; and because of this shortfall additional novel vaccine development is urgently required. However, despite the long history to develop vaccines which target blood stage parasites, a clinically approved vaccine remains elusive [4,5].

The secretory protein apical membrane antigen 1 (AMA1), first identified as an integral membrane protein of the *P. falciparum* apical complex [6], is a leading target for vaccines intended to reduce the efficiency of parasite proliferation in the patient blood. The protein is essential for merozoite invasion of erythrocytes via interacting with rhoptry neck protein 2 (RON2) which has been prior inserted in the erythrocyte cellular membrane [7–10]; and intervention measures might target this interaction. After merozoites attach to erythrocytes, the proteins RON2, RON4, and RON5 are secreted from apical organelles termed rhoptries, and form a complex in the erythrocyte cellular membrane; then AMA1, secreted from a second apical organelle termed micronemes, interacts with RON2, and confers merozoite invasion into the erythrocytes [11–14]. Because this interaction occurs on the external faces of the parasite and its target host cell, it might thereby be interfered with using specific antibodies or peptides; and thus AMA1 and its interaction with the RON complex could be a promising candidate for disease-control vaccines. However, clinical vaccine trials did not show significant reduction in parasitemias in humans, likely due to the great breadth of AMA1 surface antigen polymorphisms present in parasite field populations [15–18].

AMA1 is also expressed in sporozoites, which are the infectious stage inoculated by mosquitoes into human skin during a blood meal. Sporozoite AMA1 plays a role in the invasion of mosquito salivary glands as well as the infection of hepatocytes [19]; and might also function via interaction with the RON complex. RON2, RON4, and RON5 are expressed in sporozoites and have been shown to be crucial for invasion of mosquito salivary glands and mammalian hepatocytes, using a sporozoite stage-specific gene repressing system in *P. berghei* [20–22]. Accordingly, AMA1 could be a potential target of multifunctional vaccines; that is, reducing both sporozoite and merozoite infectivity.

Here, to establish a novel screening platform for identifying potential dual-functional vaccines or small-molecules targeting the AMA1-RON2 interaction, we utilized genome editing to generate a chimeric rodent malaria parasite (*P. berghei*) that expresses PfAMA1 in place of endogenous PbAMA1. Despite the observation that some amino acid residues which were determined to be involved in interaction with RON2 are not conserved between PfAMA1 and PbAMA1, the infectivity of chimeric merozoites and sporozoites were comparable to the *P. berghei* parental line. Therefore, we propose that our chimeric parasites could serve as a useful platform to screen novel inhibitory antibodies and reagents, including those interfering with complex formation between AMA1 and RON2. In addition, infection of chimeric blood-stage parasites in rats induced inhibitory antibodies against *P. falciparum* blood-stage proliferation in a strain-transcending manner; suggesting that structure dependent antibodies could be induced and isolated using the Pb_PfAMA1 chimeric rodent malaria parasites. To expand this utility, we also generated doubly mutated chimeric parasites expressing PfAMA1 and PfCSP; and thus complemented two leading vaccine targets for study of disease control and infection blocking.

## Results

### Comparison of the secondary structure of AMA1 across *Plasmodium* species and the effects of polymorphisms on its structure

AMA1 possesses an N-terminal signal peptide followed by domains I, II, and III, with well conserved amino acid (aa) sequences among *Plasmodium* spp. (S1A Fig). Residues located on the surface of hydrophobic troughs, shown as blue highlights in S1A Fig, are highly preserved in terms of their physicochemical properties; suggesting that the AMA1 and RON2 interaction could be crucial for host infection across *Plasmodium* spp. [10,23]. Although the overall sequence identity between PfAMA1 and PbAMA1 is 45%, a residue in PfAMA1 crucial for interaction with RON2 is not conserved in PbAMA1; specifically, a phenylalanine at aa 183. Moreover, while the C-terminus loop region in PfRON2 (PfRON2sp1) is well conserved and known to interact with AMA1, two of the four residues critical for AMA1 interaction differ in PbRON2sp1 (S1B Fig, [10,24]). Accordingly, we compared the predicted structural compatibility of the heterologous PfAMA1-PbRON2sp1 interaction with that of the homologous interactions. As indicated by the corresponding ipTM values in Fig 1A, the heterologous complex is predicted with high confidence similarly to the homologous complexes. Superimposing the left and middle models suggests that PfRON2sp1 and PbRON2sp1 can occupy the hydrophobic trough of PfAMA1 in a similar mode (Fig 1B). In addition, AlphaFold analysis suggests that R1, a known peptide which inhibits the PfAMA1-PfRON2 interaction [8], overlaps with the same binding pocket and could compete with the PfAMA1-PbRON2sp1 interaction (Fig 1C).

**Fig 1.**
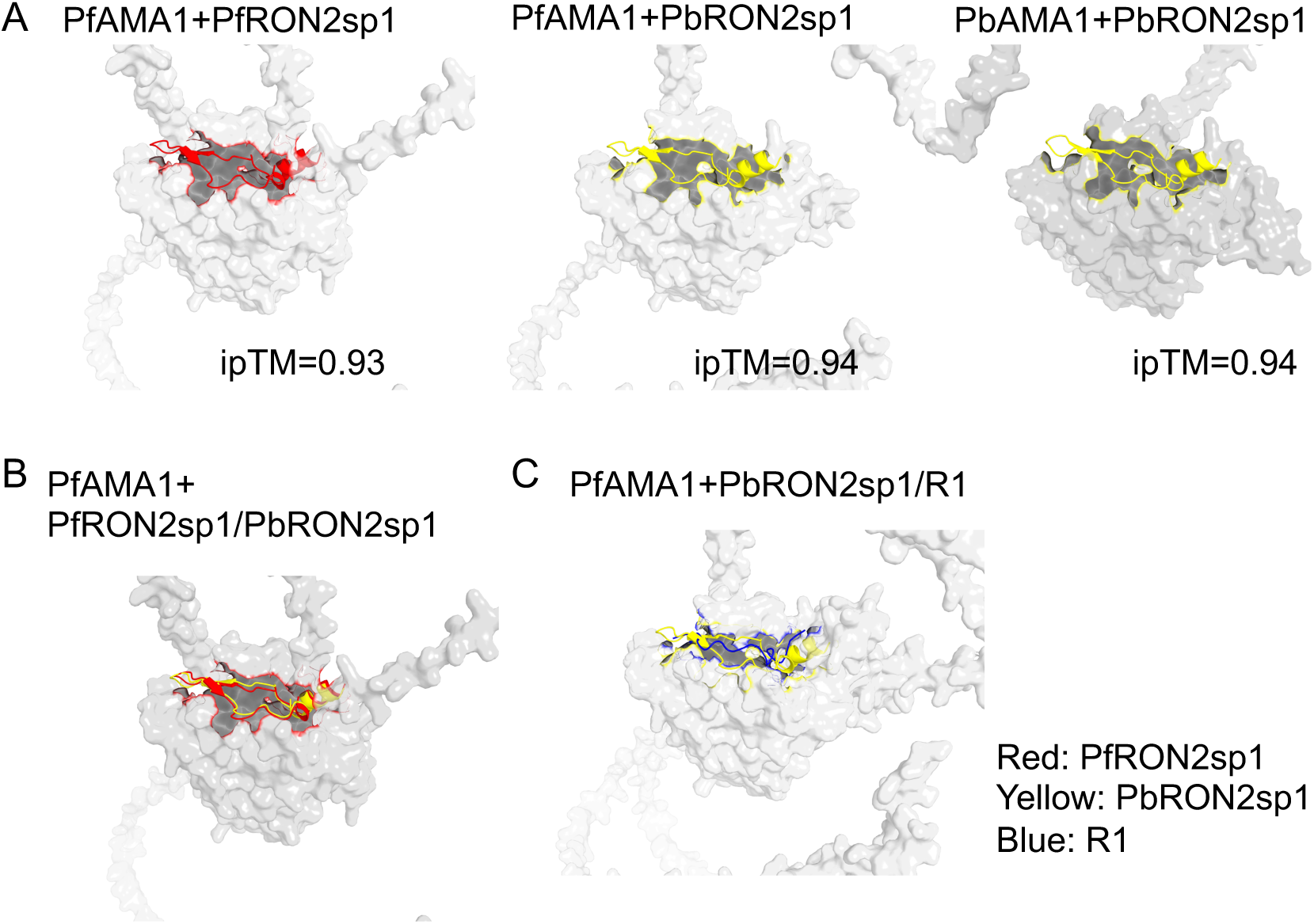
Prediction of heterogenic interaction between PfAMA1 and PbRON2. (A) Homologous and heterologous interaction between AMA1 and a known interacting peptide from the C-terminus region of RON2 (RON2sp1) is predicted by Alphafold3. The surfaces of PfAMA1 and PbAMA1 are shown in light grey and grey, respectively. PfRON2sp1 (aa 2021-2059, [10]) and corresponding PbRON2sp1 (aa 1913-1951) are shown in red and yellow cartoons. (B) Overlay image of the two predicted complexes, PfAMA1+PfRON2sp1 and PfAMA1+PbRON2sp1. Both PfRON2sp1 and PbRON2sp1 are predicted to occupy the hydrophobic trough of PfAMA1 in a similar mode. (C) Overlay image of the two predicted complexes between PfAMA1+PbRON2sp1 and PfAMA1 and the known inhibitory peptide R1 (blue). R1 peptide is predicted to occupy an overlapping region of PfAMA1 with PbRON2sp1.

### Generation of chimeric rodent malaria parasites complemented PbAMA1 with PfAMA1

To investigate whether PfAMA1 could complement the essential roles of PbAMA1 throughout the lifecycle, we generated a transgenic *Plasmodium berghei* rodent malaria parasite expressing PfAMA1 under the *pbama1* promoter at the native *pbama1* locus. The GFP-Akaluc parasite, in which GFP and Akaluc fused with a T2A skip peptide are constitutively expressed in the cytosol [25], was used as the parental line to replace the PbAMA1 coding region with PfAMA1 utilizing the CRISPR/Cas9 system (Fig 2A). The targeted locus in a cloned transgenic parasite was correctly replaced by the Pf_3D7_AMA1 coding region, demonstrated by genotyping PCR and DNA sequencing; and designated as Pb_PfAMA1 (Fig 2B). The resulting chimeric parasites were designed to retain the PbAMA1 signal peptide (aa 1-21 of PBANKA_0915000) and express Pf_3D7_AMA1 (aa 25-622 of PF3D7_1133400) (Fig 2C). Pb_PfAMA1 schizonts were enriched after in vitro culture of infected erythrocytes and PfAMA1 expression was examined by western blotting using an anti-AMA1-C antibody which recognizes a C-terminal peptide highly conserved between PfAMA1 and PbAMA1 [26], shown in Figure 2C). As shown in Fig 2D, two bands were detected by anti-AMA1-C antibodies in both Pb_PfAMA1 and Pf3D7 schizonts, at approximately 75 kDa and 64 kDa; demonstrating that PfAMA1 is expressed normally in Pb_PfAMA1 chimeric merozoites and the N-terminal shedding occurs in the chimeric parasites in the same manner as in *P. falciparum* [27]. The antibody detected PbAMA1 from GFP-Akaluc schizonts at approximately 56 kDa, corresponding to the calculated full-length PbAMA1, with a possible processed form at 18 kDa. This result confirmed that Pb_PfAMA1 chimeric merozoites expressed only PfAMA1, and no PbAMA1. By immunoelectron microscopy using anti-PfAMA1 antiserum [28], PfAMA1 was demonstrated to localize to micronemes in Pb_PfAMA1 merozoites in schizont stage (Fig 2E). The apical localization of PfAMA1 in Pb_PfAMA1 merozoites was also detected by immunofluorescence analysis using anti-AMA1-C antibodies (S2 Fig).

**Fig 2.**
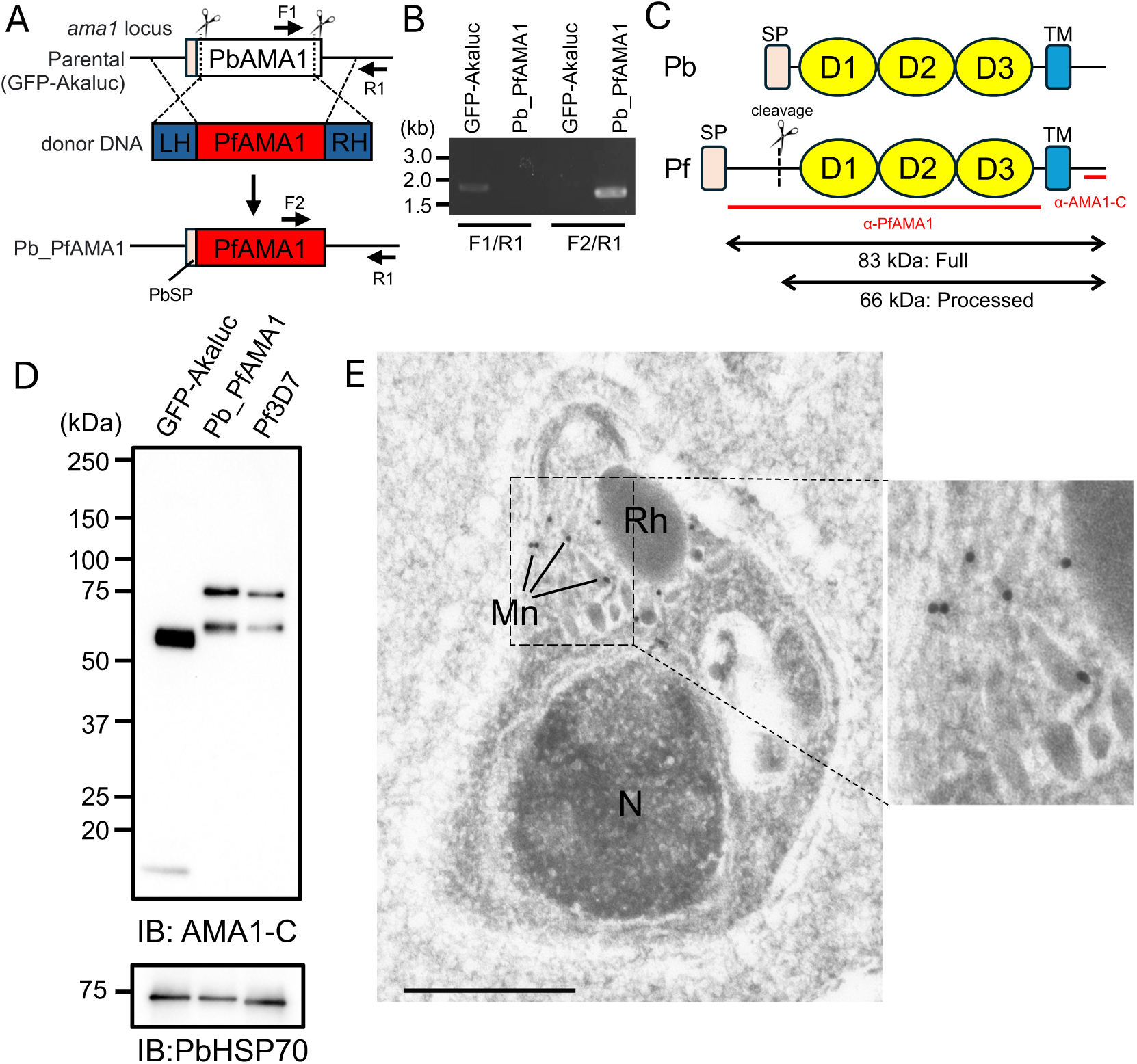
Generation of a chimeric *Plasmodium berghei* parasite expressing PfAMA1 instead of PbAMA1. (A) Schematic representation of the generation of transgenic parasites in which the PbAMA1 coding region is replaced with the PfAMA1 coding sequence using the CRISPR/Cas9 system. The entire PbAMA1 coding region except for the signal peptide was replaced by the Pf_3D7_AMA1 coding sequence by double digestion with Cas9, as indicated by scissors. The transgenic parasites express PfAMA1 under the PbAMA1 promotor, designated as Pb_PfAMA1. (B) PCR genotyping demonstrating replacement of the PbAMA1 coding region. After cloning, the transgenic parasite genotype was investigated by PCR directly using infected mouse blood as template and the primer sets indicated in (A) (F1: PbAMA1-check-F; F2: PfAMA1-check-F; R1: AMA1check-R). Using the *pfama1* specific primers (F2/R1), an expected DNA fragment at about 1.5 kbp was amplified from infected mouse blood with cloned Pb_PfAMA1 transgenic parasites. (C) Schematic secondary structures of PbAMA1 and PfAMA1. PfAMA1 (83-kDa) is processed into a 66-kDa mature form near the signal peptide (SP), while no processed site was reported in PbAMA1. Antigens used for raising antibodies, anti-PfAMA1 and anti-AMA1-C antibodies, are indicated by red bars. TM, transmembrane domain. (D) Western blotting showing that Pb_PfAMA1 schizonts express PfAMA1 instead of PbAMA1. Parental and Pb_PfAMA1 purified schizonts (1 x 10^6^) were loaded, along with the same number of Pf3D7 schizonts, and western blotting was performed using anti-AMA1-C antibodies (upper panel, 1:10,000 dilution). Two bands at approximately 75 kDa and 64 kDa were detected corresponding to full-length and processed forms in Pb_PfAMA1 and Pf schizonts. In the parental parasite line, bands at approximately 56 kDa and 18 kDa were detected due to the cross reactivity of antibodies. The same membrane was incubated with anti-PbHSP70 antiserum (1:500,000 dilution) to show protein loading. Specific bands were detected at about 70 kDa as reported [45]. (E) PfAMA1 is localized to micronemes in Pb_PfAMA1 schizonts. Immunoelectron microscopy was performed using anti PfAMA1 antibodies (1:500 dilution). Gold particles indicate PfAMA1 localization mostly in micronemes at the apical end of a merozoite. The dotted box indicates the region selected for magnification on the right. Bar, 500 nm; Mn, microneme; Rh, rhoptry; N, nucleus.

### PfAMA1 complements the role of PbAMA1 during parasite proliferation in mouse blood

It has been reported that conditional knockdown or knockout of *ama1* during the erythrocytic stage disrupted parasite proliferation, indicating that AMA1 is crucial for the invasion of erythrocytes [12,19]. To investigate whether PfAMA1 in Pb_PfAMA1 merozoites complements the roles of PbAMA1 during invasion, GFP-Akaluc or Pb_PfAMA1 infected erythrocytes were intravenously inoculated into five female BALB/c mice and their parasitemias over time were compared (Fig 3A and S3 Fig). Pb_PfAMA1 chimeric parasites proliferated as efficiently as the parental line, supporting functional complementation of PbAMA1 by PfAMA1 during blood-stage proliferation. Since it has been demonstrated that prior to invasion secreted AMA1 interacts with the rhoptry neck protein (RON) complex consisting of RON2, RON4, and RON5 [8], we next used a co-immunoprecipitation assay to investigate whether PfAMA1 could interact with the PbRON complex. Protein lysates of GFP-Akaluc- or Pb_PfAMA1- schizonts were incubated with anti-PbRON4 antibodies, followed by western blotting to detect RON4 and AMA1. PfAMA1 could be detected in the pull-down sample, although the band intensity was weaker compared to PbAMA1; supporting association of PfAMA1 with the heterologous PbRON complex but not permitting a quantitative comparison of binding affinity (Fig 3B). Taken together, these data indicate functional complementation by PfAMA1 in chimeric blood-stage parasites via its association with the PbRON complex inserted into the host erythrocytic membrane.

**Fig 3.**
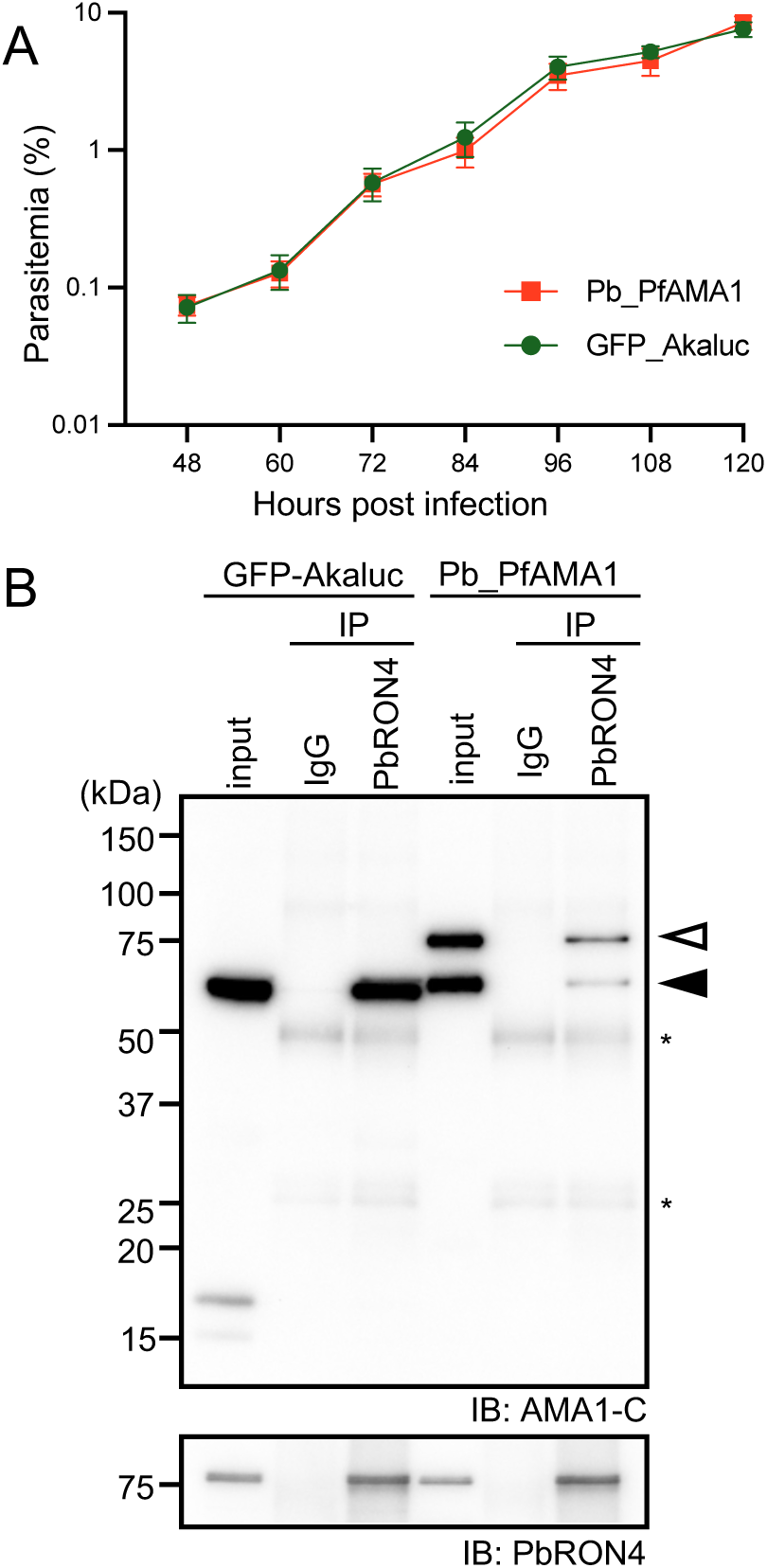
PfAMA1 complements the roles of PbAMA1 during blood stage parasite proliferation. (A) Parasitemias after inoculation of chimeric blood-stage parasites into BALB/c mice. Infected erythrocytes (1 x 10^5^) from parental (GFP-Akaluc)- or Pb_PfAMA1- parasites were intravenously inoculated into five BALB/c mice. The mean parasitemias with standard deviations are plotted, showing that PfAMA1 functions during invasion of erythrocytes by chimeric *P. berghei* merozoites. Reproducibility is shown in S3 Fig. (B) Co-immunoprecipitation assay demonstrating that PbRON4 forms a complex with PfAMA1 in chimeric parasites. Protein lysates from 1 x 10^6^ schizonts of GFP-Akaluc (parent) and Pb_PfAMA1 were incubated with anti-PbRON4 antibodies or normal IgG as control. PbAMA1 and PfAMA1 were detected by western blotting with anti-peptide antibodies recognizing the C-terminal conserved region in PfAMA1 (upper panel). Full-length and processed forms of PfAMA1 are shown by open and closed arrowheads, respectively. IgG heavy chain and light chain are indicated by asterisks. PbRON4 was detected by anti-PbRON4 antibodies to confirm the efficacy of immunoprecipitation (lower panel).

### PfAMA1 plays important roles also in chimeric sporozoites during transmission from mosquitoes to mammalian hosts

AMA1 is additionally expressed in sporozoites, which are formed within oocysts on mosquito midguts; and conditional knockout experiments indicate that the protein plays crucial roles during sporozoite invasion of salivary glands and infection of mammalian hepatocytes [19]. Western blotting demonstrated that the quantity of PfAMA1 increased upon sporozoite maturation, and its signal could be clearly detected in Pb_PfAMA1 chimeric sporozoites collected from salivary glands, similarly to PbAMA1 (Fig 4A) [29]. The localization of PfAMA1 in Pb_PfAMA1 sporozoites was detected by IFA using anti-AMA1-C antibodies together with anti-PbTRAP or anti-PbRON12 antibodies as markers for micronemes or rhoptries, respectively (Fig 4B). PfAMA1 was detected throughout the sporozoite cytosol, with higher intensity in the middle region anterior to the nucleus, like the pattern of PbTRAP. In contrast, PbRON12 localized most strongly to the apical end, which did not completely overlap with PfAMA1 signals. PfAMA1 localization was further investigated by immunoelectron microscopy using ultrathin sections of Pb_PfAMA1 sporozoites residing in salivary glands. As shown in Fig 4C, gold particles corresponding to PfAMA1 localized to sporozoite micronemes. Taken together, we demonstrated that PfAMA1 localizes to micronemes in sporozoites as well as in merozoites. To investigate whether PfAMA1 could complement PbAMA1 roles in sporozoites, sporozoite numbers were compared between parental- and Pb_PfAMA1- parasites infected mosquitoes. The average numbers of sporozoites invading salivary glands, as well as those formed inside oocysts on midguts, were not significantly different between parental line and Pb_PfAMA1 parasites; demonstrating that PfAMA1 functions sufficiently to replace the crucial roles of PbAMA1 during invasion of salivary glands (Fig 4D). The efficacy of Pb_PfAMA1 sporozoite infection of mice was examined by detection of the parasite burden in livers at 44 h after intravenous inoculation of sporozoites into five C57BL/6 mice. Pb_PfAMA1 parasite amounts in the liver were detected by red-shifted derivative luciferase activity (Akaluc)[30], which is expressed in the parasite cytosol under the *pbhsp70* promoter throughout the life stages [25], and were comparable to those of GFP-Akaluc (Fig 4E). These data support functional complementation by PfAMA1 during sporozoite infection of mouse livers. Our results demonstrated that PfAMA1 in Pb_PfAMA1 chimeric parasites plays essential roles in the two infectious stages, merozoites and sporozoites.

**Fig 4.**
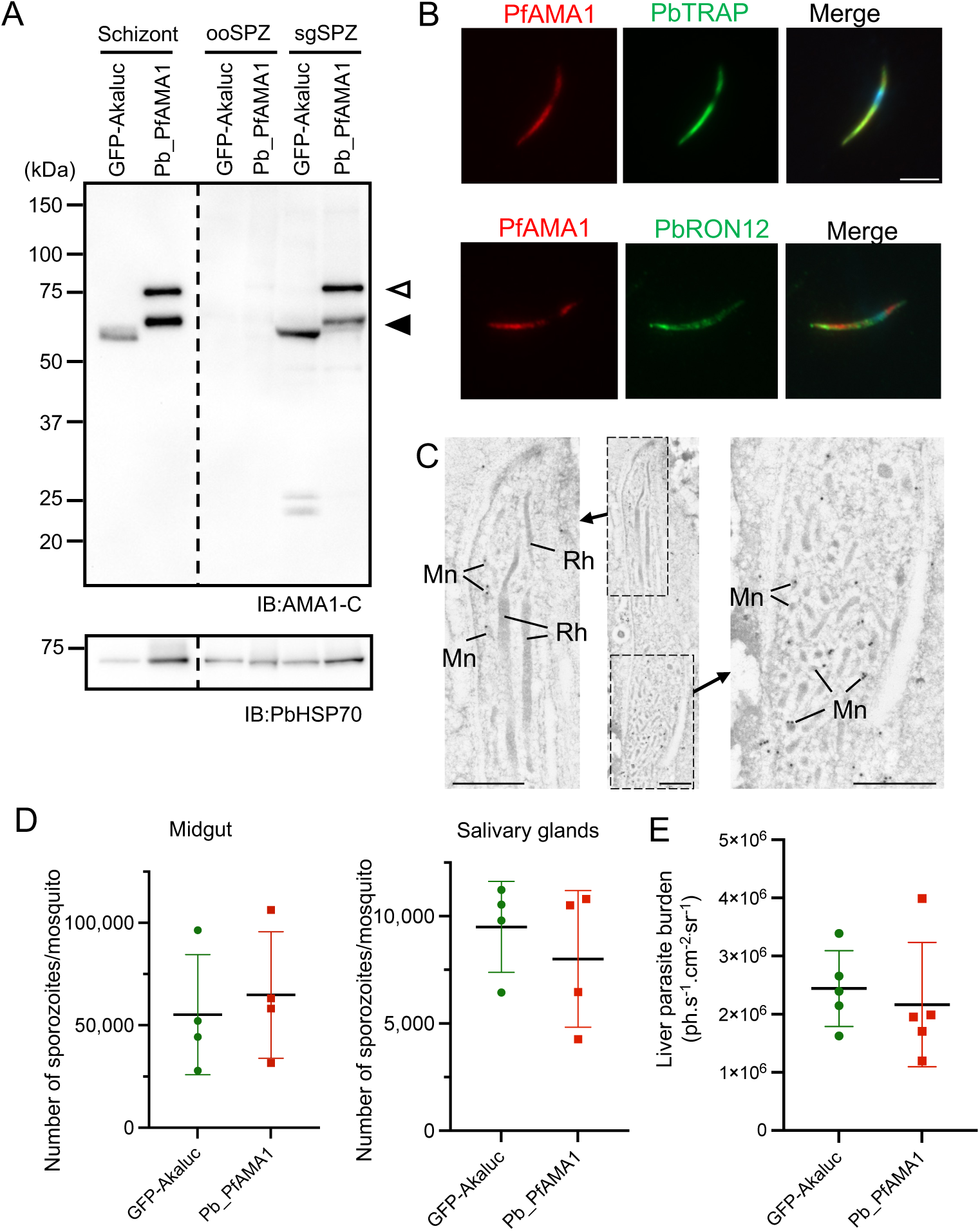
PfAMA1 in sporozoites compensate PbAMA1 roles during invasion of salivary glands and infection of mouse liver. (A) PfAMA1 expression in sporozoites collected from midguts and salivary glands of infected mosquitoes. Protein lysates from 1 x 10^5^ schizonts, oocyst-derived sporozoites (ooSPZ), or salivary glands-derived sporozoites (sgSPZ) were subjected to western blotting using anti-AMA1-C antibodies. Full-length and processed forms of PfAMA1 are shown by open and closed arrowheads (upper panels), respectively. Protein loading was examined using anti-PbHSP70 antibodies (1:5000,000) (lower panels). (B) PfAMA1 detection in salivary gland-residing Pb_PfAMA1 sporozoites examined by IFA. Acetone-fixed sporozoites were incubated with anti-AMA1-C antibodies (1:2000, shown in red) together with anti-PbTRAP antiserum (1:1000, upper panels, shown in green) or with anti-PbRON12 antiserum (1:1000, lower panels, shown in green). The merged images also show nuclei stained with Hoechst (shown in blue).

The PfAMA1 signal mostly overlaps with PbTRAP rather than with PbRON12. (C) PfAMA1 localization analysis by immunoelectron microscopy. Ultra-thin sections of Pb_PfAMA1 infected salivary glands were incubated with anti-PfAMA1 antiserum (1:500) followed by incubation with secondary antibodies conjugated to gold particles. The dashed boxes in the central diagram outline the areas presented at higher magnification in the left (most apical part) and right (middle part) panels. Gold particles corresponding to PfAMA1 are localized to micronemes. Rh, rhoptry; Mn, microneme. Bars, 500 nm. (D) Sporozoite numbers collected from midguts (left) and salivary glands (right) of parental line or Pb_PfAMA1 infected mosquitoes. Sporozoites were harvested from midguts and salivary glands at days 20 to 23 post-feeding from GFP-Akaluc (parent) or Pb_PfAMA1 infected mosquitoes. The mean numbers of sporozoites per mosquito are plotted from at least four independent experiments. Sporozoite numbers formed in oocysts and invaded salivary glands were not significantly different between GFP-Akaluc and Pb_PfAMA1. (E) Liver burdens at 44 h after sporozoite inoculation detected by luciferase activity. Sporozoites (5 x 10^3^) were inoculated intravenously into five C57BL/6 mice, and liver parasite burdens were measured by Akaluc activity at 44 h after sporozoite inoculation. Parasite loads are plotted with the means and standard errors shown by bars.

### R1 peptide targeting the PfAMA1-RON interaction reduced Pb_PfAMA1 blood-stage expansion and hepatocyte infection

The R1 peptide is known to inhibit PfAMA1- PfRON2 interaction; and consists of 20 amino acids with a 3D structure predicted to mimic the RON2 C-terminal region binding site of AMA1 [8,10,31] Addition of R1 peptide to *P. falciparum* blood stage in vitro cultures inhibits parasite growth, thus supporting the importance of the interaction between secreted AMA1 and RON2 inserted into the target cellular membrane during invasion of erythrocytes [31]. Although the amino acid sequence identity of binding sites in PbRON2 and PfRON2 is lower than 50%, the structure analysis suggests that R1 peptide may also compete with the interaction of PbRON2 and PfAMA1 (Fig 1C and S1B Fig). To investigate whether PfAMA1 functionally interacts with PbRON2 during chimeric merozoite invasion, Pb_PfAMA1 matured schizonts (1v × 10^5^) purified from in vitro culture were inoculated into mice with 200 µg R1 peptide or F5 peptide as a control [32]. Since the inhibitory effect of R1 peptide is quickly diminished in serum, the mice were inoculated again 1 h later with the same amount of R1 or F5 peptide. R1 peptide treatment inhibited Pb_PfAMA1 parasite infection at a maximum of 70% compared to the F5 peptide (Fig 5A and S4 Fig). To examine the inhibitory effect of the peptide on sporozoite infection, peptide (1 mg/ml) was added during Pb_PfAMA1 sporozoite inoculation of the hepatoma cell line HepG2. The numbers of liver stage parasites at 48 h post-sporozoite inoculation decreased approximately 90% by R1 peptide addition; however, formed LS parasites developed to similar sizes as the control (Fig 5B and S4B Fig). These results demonstrate that Pb_PfAMA1 infection is sensitive to R1 at both blood and sporozoite stages, consistent with a functional role of the PfAMA1-RON pathway during host-cell infection. The successfully infected chimeric sporozoites could develop normally regardless of R1 peptide treatment; indicating that R1 peptide mainly inhibits sporozoite productive invasion to hepatocytes, possibly via interaction with PbRON2. Accordingly, our chimeric parasites could serve as a useful platform to evaluate infection-blocking antibodies and/or reagents targeting PfAMA1 and the AMA1-RON2 interface at the merozoite and sporozoite infectious stages.

**Fig 5.**
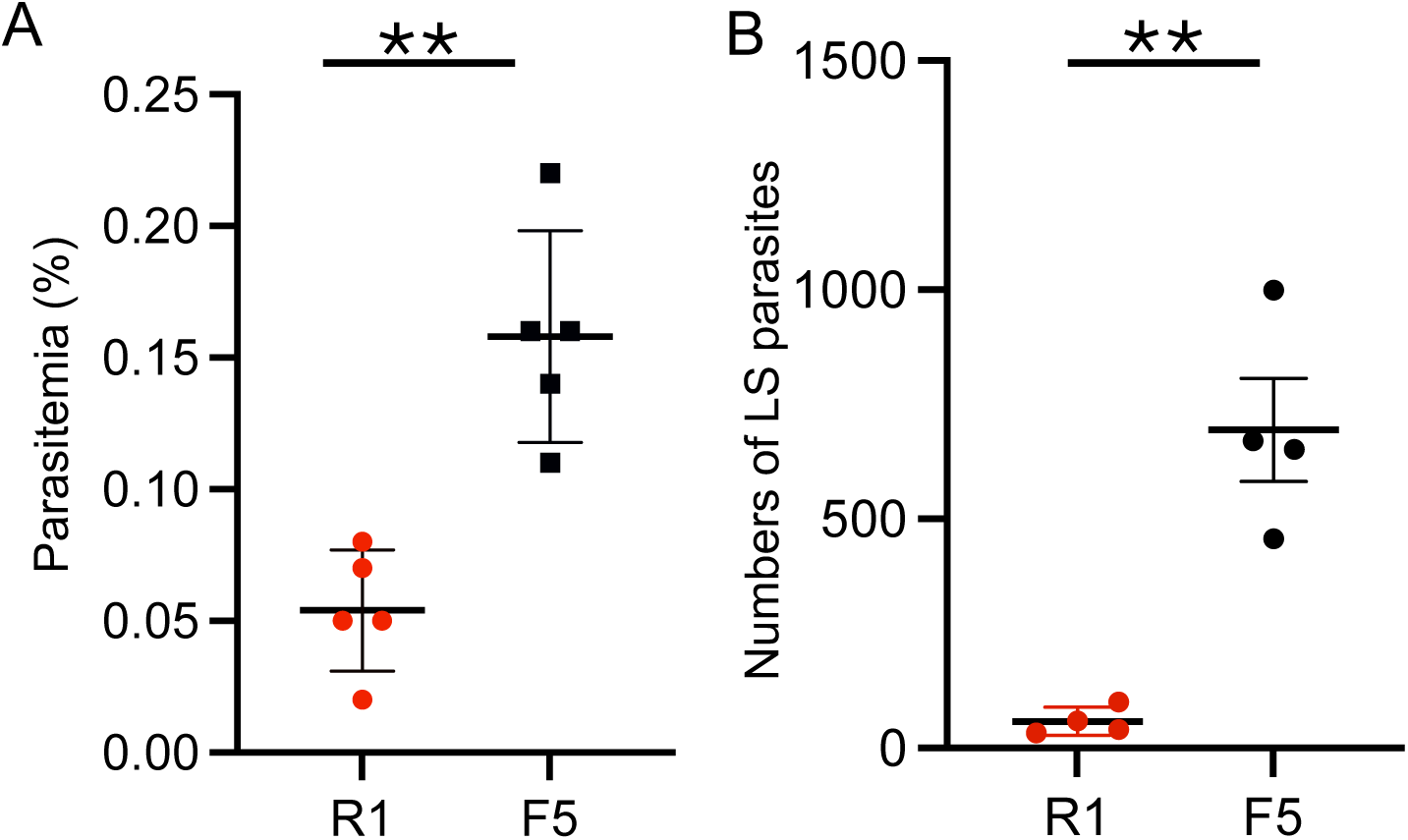
Effect of R1 peptide on Pb_PfAMA1 blood-stage proliferation and hepatocyte infection. (A) R1 peptide decreased Pb_PfAMA1 blood-stage parasite proliferation in vivo. Pb_PfAMA1 schizonts (1 × 10^5^) together with 200 µg R1 or F5 peptide were inoculated intravenously into five BALB/c mice, followed by an additional 200 µg peptide inoculation 1 h later. Parasitemias at 72 h post inoculation are plotted, showing that R1 peptide significantly decreased the subsequent parasitemia of erythrocytes in mouse blood (Unpaired t-test, **: P<0.01). The reproducibility of this experiment is shown in S4A Fig. (B) R1 peptide impaired sporozoite infection of HepG2 cells in vitro. Pb_PfAMA1 sporozoites (2 × 10^3^) were harvested from salivary glands and together with 1 mg/ ml R1 or F5 peptide were inoculated to HepG2 cells in eight-well chamber slides. The average numbers of liver stage parasites at 48 h post-sporozoite inoculation are shown as dots with means and standard errors as bars from four independent experiments. Pb_PfAMA1 sporozoite infection of hepatocytes was significantly reduced by the addition of R1 peptide (Mann-Whitney test, *: P<0.05). The areas of each liver stage parasites are shown in S4B Fig.

### Induction of infection reducing antibodies by inoculation of PfAMA1 expressing chimeric *P. berghei* blood stage parasites into rats

PfAMA1 localizes to the merozoite surface as a processed form and prior to erythrocyte invasion interacts with a C-terminal loop region of RON2. A hydrophobic trough within AMA1 is masked by its domain II until just before interacting with RON2 [10,13,33]. To investigate the hypothesis that AMA1 native structure-dependent antibodies might efficiently inhibit merozoite invasion of erythrocytes, we utilized the chimeric parasites as a live vaccine. Wister rats were immunized twice by inoculation with the Pb_PfAMA1 chimeric parasite- or GFP-Akaluc-infected erythrocytes, followed by treatment with pyrimethamine (Fig 6A). Induction of antibodies recognizing PfAMA1 in the immunized rats was confirmed by IFA on *P. falciparum* schizonts. Serum from rats immunized by Pb_PfAMA1 infection detected micronemes in *Pf* schizonts, demonstrated by co-localization with anti-PfAMA1 antiserum, preferentially to those immunized by GFP-Akaluc infection (Fig 6B). The inhibitory effects of sera raised by Pb_PfAMA1 or GFP-Akaluc infection were compared by inoculating *P. falciparum* 3D7 in vitro cultures at six times dilution. Parasite proliferation was monitored daily by dual-color DNA/RNA staining to detect parasite-infected or parasite-attached erythrocytes (Fig 6C and S5 Fig) [34]. Sera from Pb_PfAMA1-infected rats showed 97% inhibitory activity, compared to 89% inhibition by sera from GFP-Akaluc-infected rats (Fig 6D). A similar tendency was observed when sera were added to HB3B, a subclone of *P. falciparum* HB3 strain, which originated from Honduras and possesses several amino acid substitutions within AMA1 [35]; indicating inhibitory activity against both *P. falciparum* laboratory strains tested. Because sera from parental-parasite-infected rats also showed substantial inhibitory activity, possibly by cross-reactive antibodies raised by GFP-Akaluc infection, the magnitude of the PfAMA1-specific contribution cannot be determined from this experiment alone.

**Fig 6.**
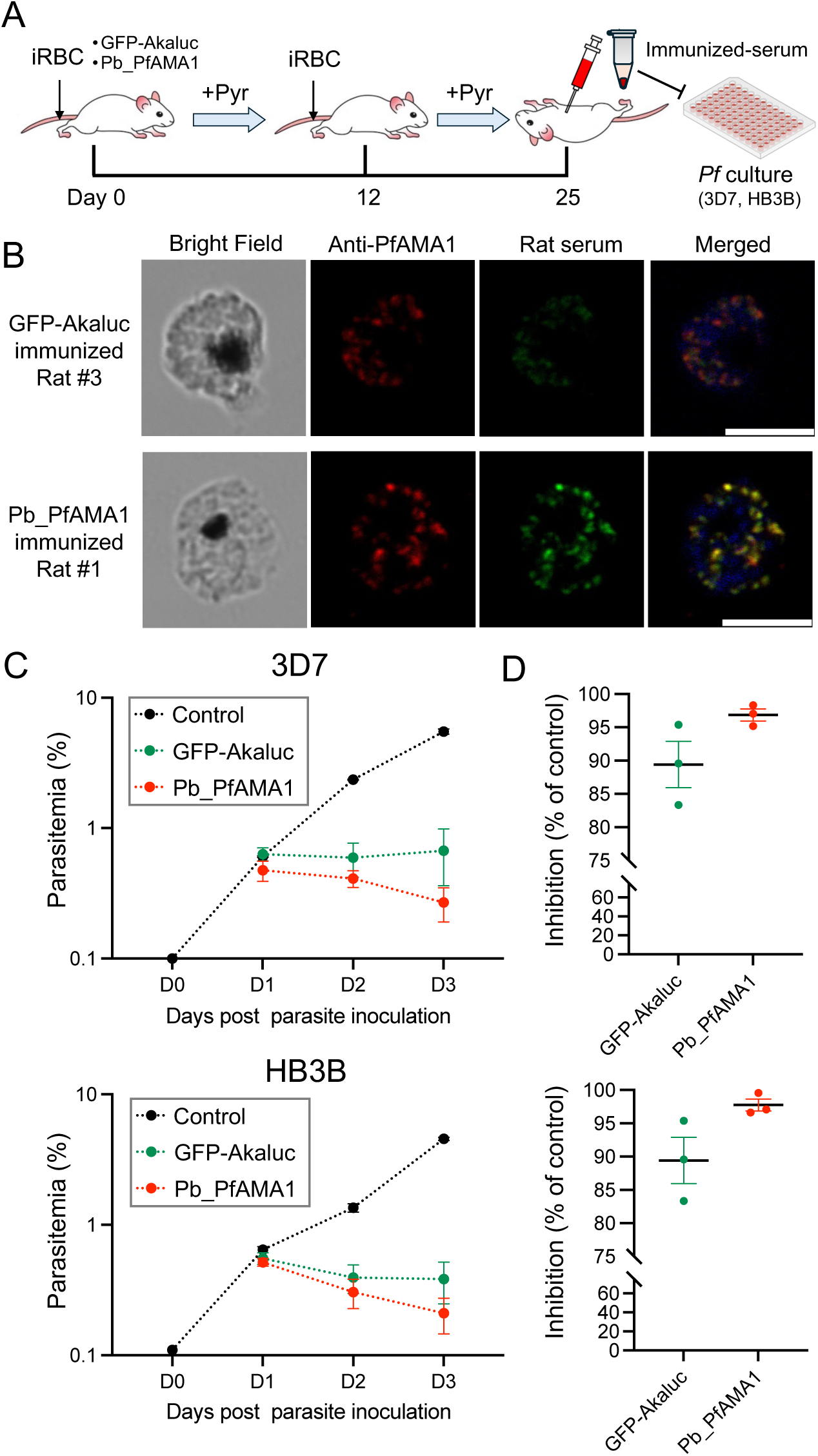
Induction of Pf merozoite infection reducing antibodies by inoculation of chimeric parasites into rats. (A) Workflow for generating immune rat serum by infection with *P. berghei* followed by chemotherapeutic cure. Rats were inoculated of 1 × 10^5^ erythrocytes infected with GFP-Akaluc or Pb_PfAMA1 parasites, followed by pyrimethamine treatment. The infection-and-treatment cycle was repeated twice, and the resulting sera were collected and used as the immunized sera for GIA. (B) Induction of PfAMA1 reactive antibodies in rats immunized by Pb_PfAMA1 parasite infection, examined by IFA. In vitro cultured *P. falciparum* 3D7 schizonts were fixed with acetone on glass slides and incubated with 1/500 diluted antisera collected from GFP-Akaluc or Pb_PfAMA1 infected rats together with anti-PfAMA1 rabbit antiserum (1:20,000). Pb_PfAMA1 immunized rat sera preferentially stained in a dotted pattern at the apical region of each merozoite (shown in green), which are co-stained with anti-PfAMA1 rabbit antiserum (red). The merged image was shown together with nuclei staining with Hoechist (blue). Bar, 5 µm. (C) Inhibitory effect of sera collected from rats immunized via Pb_PfAMA1 or GFP-Akaluc (parent) parasite infections. Tightly synchronized Pf3D7 (upper) or PfHB3B (lower) schizonts were mixed with antisera at a 1:6 dilution and the mean ratio of parasitemias plotted with standard deviations as error bars. (D) Inhibitory effects of sera on Pf3D7 (upper) and PfHB3B (lower) parasite proliferation. Inhibition levels by individual rat sera were calculated based on day 3 (D3) parasitemias compared with the control (no serum added). Data are presented as individual dots, with horizontal bars indicating the mean and standard deviation.

### Generation of doubly replaced chimeric parasites expressing PfAMA1 and PfCSP, to extend the chimeric model to two *P. falciparum* antigens

AMA1 is expressed in both merozoite and sporozoite infectious stages, and could serve as an ideal target for dual-purpose vaccines aiming to block both infection and disease progression; whereas CSP is a target solely for sporozoite liver-stage infection. To address the limitations of current vaccines, such as the incomplete sporozoite neutralization by RTS,S, novel multi-target vaccines incorporating both CSP and AMA1 are essential to interrupt the *Plasmodium* life cycle at multiple stages. Furthermore, anti-AMA1 antibodies, particularly targeting the AMA1-RON2 interaction, could suppress blood-stage proliferation of parasites that escape initial sporozoite-blocking immunity, thereby overcoming the suboptimal efficacy of existing infection-reducing vaccines. Toward this goal, we generated chimeric parasites which express PfCSP via replacement of PbCSP and confirmed that, as with the reported PfCSP-replacement transgenic line [36], no significant impairment was observed in sporogony or sporozoite infectivity. For a future platform to develop dual functional vaccines, we generated dual chimeric rodent malaria parasites expressing both PfCSP and PfAMA1, by applying the same AMA1 replacement strategy on the Pb_PfCSP parasites as a parental line. After confirming the correct integration of the PfAMA1 coding sequence by genomic PCR (S6 Fig), the cloned chimeric parasites were designated as Pb_PfCSP+PfAMA1. The influence of dual mutation on sporozoite invasion of salivary glands and infection of mouse liver was evaluated using Pb_PfCSP+PfAMA1 chimeric parasites. The numbers of sporozoites formed in oocysts and invaded salivary glands were comparable to Pb_PfCSP sporozoites (Fig 7A). In addition, the parasite burdens in mouse livers at 44 h after the inoculation of 2000 Pb_PfCSP+PfAMA1 sporozoites were not significantly different compared to mice inoculated with Pb_PfCSP sporozoites (Fig 7B). Therefore, the Pb_PfCSP+PfAMA1 chimeric parasite line provides a feasible model for future evaluation of multistage and multi-antigen interventions.

**Fig 7.**
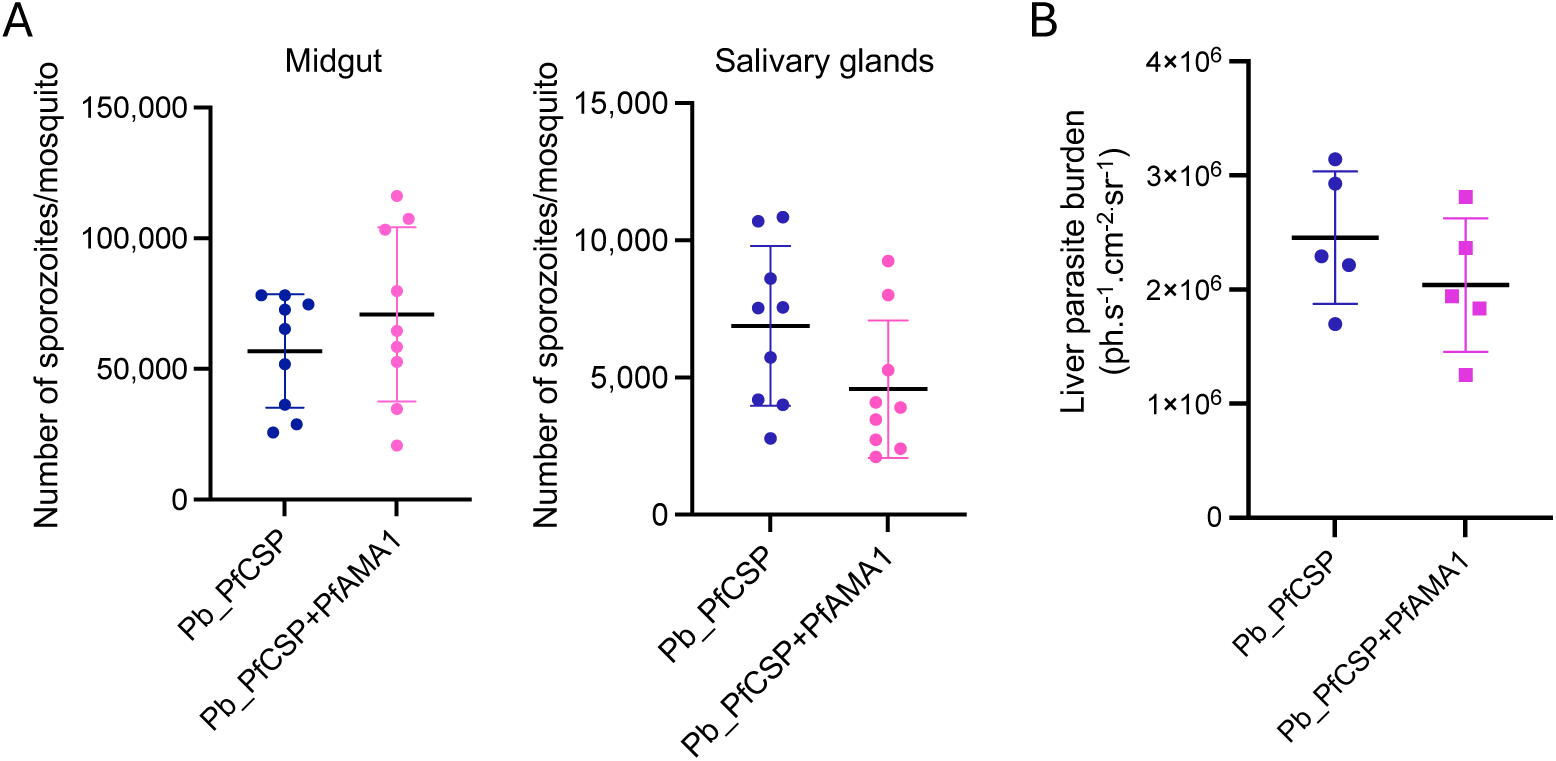
Generation of double replacement chimeric parasites expressing PfAMA1 and PfCSP. (A) Sporozoite numbers collected from midguts (left) and salivary glands (right) of Pb_PfCSP line or Pb_PfCSP+PfAMA1 infected mosquitoes. Sporozoites were harvested from midguts and salivary glands at days 20 to 23 post-feeding from chimeric parasite-infected mosquitoes. The mean numbers of sporozoites per mosquito are plotted from at least nine independent experiments. Sporozoite numbers formed in oocysts and invaded salivary glands are not significantly different between Pb_PfCSP and Pb_PfCSP+PfAMA1. (B) Pb_PfCSP+PfAMA1 liver burdens at 44 h after sporozoite inoculation detected by luciferase activity. Sporozoites (5 x 10^3^) were intravenously inoculated into five C57BL/6 mice and liver parasite burdens were measured by Akaluc activity at 44 h after sporozoite inoculation. Parasite loads are plotted with the means and standard errors shown by bars.

## Discussion

### PfAMA1 compensates for PbAMA1 essential functions during parasite invasion of target cells

The PbAMA1 coding region was successfully replaced by PfAMA1 within the *Plasmodium berghei* genome; and the resulting Pb_PfAMA1 chimeric parasites infected erythrocytes and mouse liver as efficiently as the parental *P. berghei* parasite line. Taken together with the report that PbAMA1 is essential for blood stage parasite proliferation and invasion of salivary glands and hepatocytes [12,19], PfAMA1 successfully replaces the crucial roles of PbAMA1 in two infectious stages, sporozoite and merozoite, of Pb_PfAMA1 chimeric parasites. Although amino acid identity between PbAMA1 and PfAMA1 is about 45%, AlphaFold 3 predicts structurally compatible interactions between PfAMA1 and the PbRON2 C-terminal loop, PbRON2sp1, as well as within the homologous complex (Fig 1). This prediction is supported by a co-immunoprecipitation assay demonstrating that anti-PbRON4 antibodies precipitate PfAMA1 together with the PbRON complex (Fig 3B); however, the relative band intensities do not permit a quantitative comparison of binding affinity. It should be noted that co-immunoprecipitation assays do not account for the fact that in schizonts AMA1 and RON2 are present in different apical organelle compartments, micronemes versus rhoptries, and further investigations such as cryo-electron microscopy might be required to determine the functional interaction of PfAMA1 and PbRON2 after their secretion upon Pb_PfAMA1 merozoite invasion of erythrocytes.

### Evidence consistent with functional interaction of PfAMA1 with the PbRON pathway during Pb_PfAMA1 blood-stage infection

The C-terminal region of PfRON2 (amino acid 2027-2047) interacts with a hydrophobic trough on the surface of PfAMA1 [10,13]. R1 peptide was selected from several synthesized peptides derived from the C-terminal region of PfRON2 which interact with the hydrophobic trough of PfAMA1 and exhibit invasion inhibition efficacy [31]. Despite a sequence identity of 25%, the IC_50_ of R1 peptide in *P. falciparum* merozoite invasion of erythrocytes was 4 µM, due to its 3D structural mimicry of the RON2 binding site and high affinity interaction with the PfAMA1 hydrophobic pocket [31]. In this study, we attempted to inhibit Pb_PfAMA1 chimeric merozoite invasion of mouse erythrocytes using the R1 peptide in vivo, as *P. berghei* blood-stage parasites cannot efficiently proliferate in vitro. Since the R1 is unstable in serum [37], we treated mice with 200 µg of R1 peptide twice, once during the inoculation of enriched-mature schizonts and a second dose 1 h post-inoculation. R1 peptide reduced parasite proliferation in mouse blood to approximately 30% of a control using scrambled peptide (Fig 5 and S4 Fig). Although the R1 peptide inhibitory effect on Pb_PfAMA1 was lower than that on *P. falciparum* merozoites, this experiment supports that the heterologous binding of PfAMA1–PbRON2 contributes to merozoite invasion of erythrocytes.

### Interaction inhibiting reagents as novel malaria intervention development

PbAMA1 conditional knockdown demonstrated that sporozoite AMA1 is important for both the invasion of mosquito salivary glands and mammalian hepatocytes [19]. However, its mode of action remains largely unknown, including whether AMA1 functions through interaction with RON2. In addition, its precise localization within sporozoites has not been clearly visualized; and here for the first time we demonstrated a micronemal localization by immunoelectron microscopy using our chimeric rodent malaria parasites (Fig 4C). Evaluation of this interaction using native *P. falciparum* sporozoites has been hindered by substantial technical limitations, particularly the difficulty of collecting sufficient infectious *P. falciparum* sporozoites from infected mosquitoes and the lack of an efficient in vitro infection system. For example, while the R1 peptide at 1 mg/ml was reported to reduce *P. falciparum* sporozoite invasion of cultured HC-04 hepatocytes by approximately 59% relative to the DMSO control, this inhibition was attributed to effects on both cell traversal and invasion [38]. Notably, in the current study, R1 peptide at the same concentration inhibited approximately 90% of Pb_PfAMA1 sporozoite infection of HepG2 cells, determined by the numbers of matured liver stages, corresponding to sporozoite infection accompanied by parasitophorous vacuole membrane formation. Considering that successfully invaded chimeric sporozoites develop normally inside cells, our results indicate that the AMA1–RON2 interaction plays a crucial role also during productive sporozoite infection of hepatocytes. This demonstration was achieved by exploiting our chimeric parasite system, which overcomes the difficulty of a native *P. falciparum* sporozoite in vitro infection assay. Indeed, the efficacy of *P. berghei* sporozoite infection of HepG2 cells is significantly higher and more robust than those of *P. falciparum* using any type of hepatocyte. Consequently, we propose that our chimeric rodent malaria sporozoites provide a useful and safer tool to evaluate reagents or antibodies targeting PfAMA1, and the AMA1–RON2 interaction during sporozoite infection of the liver.

### Structure based vaccine development/ screening

PfAMA1 has been a leading candidate to develop disease-control vaccines; however, the attempts to immunize humans with recombinant PfAMA1 protein did not sufficiently protect them from parasite proliferation in the blood, mainly due to the high polymorphisms in AMA1 surface antigens across field-isolated parasites [15,17]. Recently, cryo-electron microscopy revealed the precise structure of the protein complex consisting of AMA1, RON2, RON4, and RON5 at the moving junction and demonstrated that AMA1 structure was modified through its interaction with RON2 [14]. Our Pb_PfAMA1 chimeric blood-stage parasites also likely express PfAMA1 on the merozoite surface and subsequently form a complex with RON2 inserted into the erythrocyte membrane. Therefore, infection of Pb_PfAMA1 parasites may facilitate the induction of antibodies which recognize the native structure of PfAMA1 in complex with the RON invasion apparatus in infected rats. In this study, two immunizations of Wistar rats with 1 × 10^5^ Pb_PfAMA1 infected erythrocytes induced PfAMA1-reactive antibodies, and sera showed inhibitory activity against two *P. falciparum* laboratory strains tested; however, the substantial inhibition observed using the parental-parasite immune sera limits an attribution of this activity specifically to PfAMA1. Further screening of PfAMA1-reactive monoclonal antibodies, isolated from infected rodents, could lead to the development of novel interventions with enhanced inhibitory activity across both merozoite and sporozoite stages. In this study, we successfully generated a Pb_PfCSP+PfAMA1 chimeric rodent malaria parasite with comparable infectivity to liver and erythrocytes in mice. By applying this chimeric parasite, antibodies directed against two major promising targets during sporozoite infection could be evaluated in the mouse model. It is also possible that sporozoite infection-reducing anti-AMA1 antibodies could attenuate parasite proliferation in the blood-stage, even after some sporozoites escaped from infection blocking antibodies. In addition, the model chimeric parasites could be further utilized as live vaccines to investigate the cellular immune response against PfCSP and PfAMA1 in the mouse model.

## Methods

### Parasites, mice, and mosquitoes

The experimental protocols which involved rodents were reviewed and approved by the Animal Experiment Committee of the Institute of Science, Tokyo (A2026-088A, A2026-075A, A2026-087C). Mice were maintained under controlled environmental conditions with a 12-hour light/dark cycle and ambient temperature. *Anopheles stephensi* SDA500 mosquitoes were provided with a 5% (w/v) sucrose solution and maintained at 25°C. All transgenic parasites were derived from the *Plasmodium berghei* ANKA strain and were maintained in female ICR mice, aged 4 to 6 weeks.

### Plasmodium falciparum in vitro culture

The 3D7 (MRA-102) and HB3B (MRA-1227) laboratory strains of *P. falciparum* were obtained from the MR4 repository (BEI Resources, Manassas, VA, USA). The parasites were cultured in human type O erythrocytes obtained from the Japanese Red Cross Tokyo Blood Center at 2% hematocrit in a complete medium, which consisted of RPMI-1640 medium containing 2.5% human serum (obtained from the Japanese Red Cross Tokyo Blood Center), 0.25% AlbuMAX II (Life Technologies, Carlsbad, CA, USA), 25 mM HEPES, 0.225% sodium bicarbonate, 0.38 mM hypoxanthine, and 10 μg/ml gentamicin. Parasite cultures were maintained under low-oxygen conditions consisting of 90% N₂, 5% CO₂, and 5% O₂, as described [39]. Parasitemias were measured using a dual-color flow cytometric staining method, as described [34]. *P. falciparum* infected erythrocytes were diluted 10-fold with D-PBS containing 5 µg/mL dihydroethidium (DHE) (Sigma) and 8 µM Hoechst 33342 (Thermo Scientific), and incubated at room temperature for 20 min. The stained samples were then further diluted 10-fold with ice-cold D-PBS. In viable cells, DHE is converted to ethidium, which stains both DNA and RNA. Parasite developmental stages were therefore distinguished on the basis of Hoechst 33342 and ethidium fluorescence intensities, reflecting DNA content and transcriptional activity, respectively. Samples were analyzed using a MACSQuant Analyzer VYB (Miltenyi Biotec, Bergisch Gladbach, Germany), with a violet laser (405-nm excitation) used for Hoechst 33342 and a blue laser (488 nm excitation) used for DHE. Approximately 200,000 events were acquired for each sample. Data were analyzed using MACSQuantify software, with debris and doublets excluded before gating of infected erythrocytes and parasite developmental stages. Parasitemia (%) was calculated as the sum of ring-, trophozoite-, and schizont-stage events divided by the total number of events remaining after exclusion of debris and doublets.

### Generation of transgenic parasites

To generate transgenic *P. berghei* expressing PfAMA1, the *Pb*AMA1 coding region in the GFP-Akaluc parental line, which constitutively expresses GFP and Akaluc separated by a T2A skip peptide (Sekine et al., 2026), was replaced with the *Pf*AMA1 coding region using the CRISPR/Cas9 system, with some modifications to a described method [40]. The *Pf*AMA1 coding region lacked its native signal peptide, thereby retaining the endogenous *Pb*AMA1 signal peptide. The donor DNA was generated by overlap extension PCR to fuse the left and right homology arms and PfAMA1-coding DNA. The left and right homology arms were amplified from *P. berghei* genomic DNA using PbAMA1-LH-F/ PbAMA1-LH-R and PbAMA1-RH-F/ PbAMA1-RH-R, respectively. The PfAMA1-coding DNA was amplified from *P. falciparum* 3D7 genomic DNA using PfAMA1-F/ PfAMA1-R. These three PCR fragments were fused by overlap extension PCR, and the fused DNA fragments were used as linear donor DNA for homology-directed repair. The pCas9/sgRNA2 plasmid, which contains Cas9 and two distinct sgRNA expression cassettes driven by the *P. falciparum* U6 and *P. yoelii* U6 promoters [41], was used for replacement of the *PbAMA1* gene. Potential guide RNA sequences (20 bp) were identified using the ChopChop program (https://chopchop.cbu.uib.no), selecting those with high efficiency and no mismatches to minimize off-target effects. Complementary oligonucleotides were annealed and ligated into the digested Cas9 plasmids with BsmBI or BsaI as described [40]. The resulting plasmids were used for genome editing. Ten micrograms of linear donor DNA and 10 µg of Cas9/sgRNA-expressing plasmid were mixed in P3 primary cell solution (Lonza, Basel, Switzerland) and used immediately to transfect roughly 10^7^ tightly synchronized GFP-Akaluc line or Pb_PfCSP line schizonts, in which the endogenous *PbCSP* coding region had been replaced with the *PfCSP* coding region derived from the *P. falciparum* 3D7 strain, using FI-115 program and Nucleofector-4D (Lonza). The transfected parasites were intravenously injected into mice, and the mice were treated with pyrimethamine for 5 days, followed by drug withdrawal. After confirming the emergence of parasites in the peripheral blood, PCR-based genotyping was performed with infected blood and specific primers. Transgenic parasite clones were obtained by the limiting dilution method, as described [40].

### PCR genotyping

Genotyping PCR was performed using mouse blood containing parasites that emerged after drug withdrawal as described [42]. Tail blood was diluted 5-fold with distilled water and used as a PCR template. Primers were designed to bind specifically to the respective target gene regions but not to the donor DNA. PCR amplification was conducted using Takara Ex Premier DNA Polymerase. The PCR products were resolved by electrophoresis on a 1.0% agarose/TAE gel, stained with GelGreen (Fujifilm Wako, Japan), and visualized. The amplified fragments were then subjected to direct Sanger sequencing to confirm the successful replacement.

### AlphaFold-based structural analysis

Amino acid sequences of PfAMA1 (PF3D7_1133400), PbAMA1 (PBANKA_0915000), PfRON2 (PF3D7_1452000), and PbRON2 (PBANKA_1315700) were retrieved from PlasmoDB (release 68, https://plasmodb.org), and aa 2021-2059 of PfRON2 and aa 1913-1951 of PbRON2 were defined as PfRON2sp1 and PbRON2sp1, respectively. Complex structures of each PfAMA1 with PfRON2sp1, PbRON2sp1, or R1, and of PbAMA1 with PbRON2sp1, were predicted using AlphaFold3 (https://alphafoldserver.com/). Complex-structure visualization and structural superimpositions were performed using PyMOL. The confidence of each predicted complex was indicated by ipTM values [43].

### Schizont preparation of Pb and Pf parasites

For *P. berghei* schizont enrichment, blood samples from ICR mice infected with GFP-Akaluc or Pb_PfAMA1 parasite were collected when parasitemias exceeded 1%. The infected erythrocytes were cultured in medium containing RPMI 1640 (FUJIFILM Wako Pure Chemical Corporation, Osaka, Japan), 20% fetal calf serum (FCS), and 1% penicillin/streptomycin for 16 to 20 h at 37°C. Schizonts were enriched via a density gradient method using Optiprep solution containing 0.85% (w/v) NaCl and 10 mM Tricine-NaOH, pH 7.4 (Serumwerk Bernburg AG, Bernburg, Germany). For western blotting, purified schizonts were hemolyzed in lysis buffer (containing NH_4_Cl 80.23 g/L, KHCO_3_ 10.01 g/L, EDTA-2NA 3.72 g) with protease inhibitor cocktail (P8340, Sigma-Aldrich, Japan LLC) and then solubilized in 1.5x SDS-PAGE with 5% 2-mercaptoethanol (Nacalai Tesque, Kyoto, Japan). Mature *Plasmodium falciparum* schizonts were enriched using Percoll density gradient centrifugation as described [25]. Briefly, synchronized parasites cultures with matured schizonts were layered onto 40% to 70% Percoll density gradient solution (GE Healthcare Life Sciences) and centrifuged at 2200 × *g* for 22 min at 24°C to separate schizont-infected erythrocytes from ring stage parasites. Enriched schizonts were collected and washed with PBS, then subsequently used for western blotting and growth inhibition assays.

### Sporozoite collection from mosquitoes

To initiate infection, 4-week-old female ICR mice were intravenously injected with parental parasite line GFP-Akaluc, chimeric parasite lines Pb_PfAMA1, Pb_PfCSP, and Pb_PfCSP+PfAMA1 parasites. After 5 days, infected mice were blood-fed to adult female *Anopheles stephensi* mosquitoes and fully engorged mosquitoes were selected and kept at 20°C until dissection. After 9 to 10 days post-feeding, the numbers and prevalence of oocysts were determined. Midguts or salivary glands were harvested at 20 to 22 days post-feeding and ground to release sporozoites for counting or infection analyses.

### Western blotting

Protein homogenates were obtained from enriched schizonts after *in vitro* culture of infected mouse blood as described above, as well as oocyst-derived sporozoites, and salivary gland sporozoites harvested at 20 to 22 days post-feeding. The homogenates were solubilized in SDS-PAGE loading buffer under reducing conditions (5% 2-mercaptoethanol) and separated by electrophoresis using a specified number of parasites on 5–20% polyacrylamide gradient gels (ATTO, Tokyo, Japan). Proteins were transferred to polyvinylidene difluoride (PVDF) membranes. Membranes were blocked in 4% skimmed milk overnight and subsequently probed with primary antibodies at the following dilutions: rat anti-AMA1-C antibody (1:10,000 and 1:5000, MRA-897A), and rabbit anti-PbHSP70 antiserum (1:500,000, [22]) for 90 minutes at room temperature under shaking conditions. The samples were then washed in 1xTBST and secondary antibody incubation with horseradish peroxidase (HRP)-conjugated Goat anti-rat IgG (H+L) (112-035-143, 1:10,000, Jackson ImmunoResearch Laboratories, PA, USA) or HRP-conjugated goat anti-rabbit IgG (H+L) (111-035-144, 1: 10,000 dilution, Jackson ImmunoResearch Laboratories) for 1 hr. Proteins were detected using Immobilon Western Chemiluminescent HRP Substrate (Millipore, MA, USA) and imaged using the Invitrogen iBright CL1500 Imaging System (Thermo Fisher Scientific, Waltham, MA, USA).

### Indirect immunofluorescent assay

Enriched schizonts and sporozoites collected from salivary glands at 20 to 22 days post-feeding were seeded onto ten-well slides, air dried, and immediately fixed in 4% paraformaldehyde (PFA) for 20 minutes and permeabilized with acetone for 3 minutes. The slides were blocked with Blocking One Histo (NacalaiTesque, Kyoto, Japan) at room temperature for 15 to 30 minutes and then incubated with primary antibodies at the following dilutions: rat anti-AMA1-C (1:2000, MRA-897A), rabbit anti-PbTRAP (1:1000, [44]), rabbit anti-PbRON12 antiserum (1:1000, [45]), rabbit anti-PbCSP repeat antiserum (1:2000, [46]) and mouse anti-PfCSP monoclonal antibody (1:2000, MRA-183A) in PBST containing 5% Blocking One Histo at 4°C overnight. This was followed by secondary antibody incubation for 1 h at room temperature with goat anti-rat IgG (H+L) cross-adsorbed secondary antibody, Alexa Fluor Plus 594, 1:500 (Thermo Fisher Scientific); goat anti-rabbit IgG (H+L), cross-adsorbed secondary antibody, Alexa Fluor Plus 488, 1:500 (Thermo Fisher Scientific); goat anti-rabbit IgG (H+L), cross-adsorbed secondary antibody, Alexa Fluor plus 594 (Thermo Fisher Scientific); and goat anti-mouse IgG (H+L), cross-adsorbed secondary antibody, Alexa Fluor Plus 488, 1:500 (Thermo Fisher Scientific). Nuclei were stained with 10 μg/ml Hoechst 33342 (Thermo Fisher Scientific). Slides were mounted with ProLong Glass Antifade Mountant (Thermo Fisher Scientific) and images taken with an inverted fluorescence microscope (Axio Observer Z1; Carl Zeiss, Oberkochen, Germany*. Plasmodium falciparum* 3D7 cultures of synchronized schizonts were purified by Percoll density gradient centrifugation as described above. The schizont-rich parasites were seeded on 10-spot glass slides, air-dried, and fixed with acetone at 4°C for 3 min. Slides were blocked with Blocking One Histo at room temperature for 10 min and then incubated with primary antibodies (rabbit anti-PfAMA1 antiserum, 1:500; immunized rat sera prepared as described above, 1:5000) in PBS containing 5% Blocking One Histo for overnight at 4°C. After washing with PBS, the slides were incubated with secondary antibodies at 500 times dilution (Goat anti-rat IgG (H+L), Cross-adsorbed Secondary Antibody, Alexa Fluor Plus 594 and Goat anti-rabbit IgG (H+L), Cross-adsorbed Secondary Antibody, Alexa Fluor plus 488) for 1 h at room temperature with 10 μg/ml Hoechst 33342 (Thermo Fisher Scientific) for nuclei staining. Slides were mounted with ProLong Glass Antifade Mountant and images were taken with a Stralis 5 confocal microscopy (Leica, Wetzlar, Germany), and following data processing was carried out using Leica Application Suite X software.

### Immunoelectron microscopy

Schizonts of parental parasite line GFP-Akaluc or chimeric parasite Pb_PfAMA1 were purified by density gradient centrifugation as described above. Infected salivary glands were dissected at day 21 post-feeding. These samples were fixed for 30 min in 1% paraformaldehyde, 0.2% glutaraldehyde, 0.1M HEPES buffer, dehydrated, and embedded in LR-White embedding media (Polysciences, PA, USA). Ultrathin sections were blocked in a blocking buffer containing PBS with 5% skimmed milk and 0.01% Tween 20 (PBS-MT). This was followed by overnight incubation with rabbit anti-PfAMA1 antiserum (1:500 dilution, [28]) at 4°C. The sections were then washed with PBS containing 0.4% Block Ace (Yukijirushi, Sapporo, Japan) and 0.01% Tween 20 (PBS-BT) after which they were incubated for 90 min at 37°C with goat anti-rabbit IgG conjugated to 15 nm of gold particles (BBI Solutions) in PBS-MT (1:40 dilution), rinsed in PBS-BT and distilled water. The grids were then stained with 2% uranyl acetate in 50% methanol and lead citrate. A transmission electron microscope was used to examine the samples (JEM-1230; JEOL, Tokyo, Japan).

### Parasite proliferation assay

Cryopreserved parasites of the parental parasite line GFP-Akaluc or the chimeric parasite line Pb_PfAMA1 were injected intraperitoneally into female ICR mice. Once parasitemias reached 0.2 to 0.5%, 1×10^5^ parasites were injected intravenously into donor 5-week-old female BALB/c mice (n=5 per group). Parasitemias of inoculated mice were measured every 12 h until 120 h after injection, by flow cytometry using MACSQuant VYB (Miltenyi Biotec GmBH, Bergisch Gladbach, Germany) and a 488nm blue laser to detect the ratio of parasite-derived GFP positive erythrocytes.

### Co-immunoprecipitation assay

Co-immunoprecipitation assays were used to detect interactions between PfAMA1 and the *P. berghei* RON complex and were performed as described, with some modifications [20]. Schizont pellets of GFP-Akaluc or Pb_PfAMA1 parasites were suspended on ice in lysis buffer consisting of 0.5% CHAPS, 0.5 mM EDTA, 0.5% protease inhibitor cocktail (539134, Millipore), and 1 mM phenylmethylsulfonyl fluoride (#8553, CST) in 1× D-PBS. Genomic DNA was sheared by passing the lysates through a 27-gauge needle. After incubation on ice for 30 min, the lysates were centrifuged to remove insoluble material, and the resulting supernatants were collected as clarified lysates. Normal rabbit IgG or purified IgG from rabbit anti-PbRON4 antiserum [22] was added at 2 mg to a clarified lysate corresponding to 1 × 10⁷ schizonts, followed by incubation for 2 h at 4°C with gentle rotation. Subsequently, 20 µl of Protein G Dynabeads (Veritas) was added, and the samples were incubated for an additional 1 h at 4°C to capture the antibody-bound protein complexes. The beads were washed sequentially once with wash buffer composed of 50 mM Tris-HCl, 0.15 M NaCl, 1 mM EDTA, and 0.5% bovine serum albumin; twice with the same wash buffer without bovine serum albumin; and finally, once with low-salt wash buffer composed of 50 mM Tris-HCl, 0.05 M NaCl, and 1 mM EDTA. Immunoprecipitated proteins were eluted by overnight incubation in 1.5 × SDS sample buffer containing 5% 2-mercaptoethanol. The samples were then heated at 95°C for 5 min and subjected to western blot analysis.

### Sporozoite infectivity to mice

Salivary gland sporozoites were collected from mosquitoes infected with GFP-Akaluc or chimeric parasite lines Pb_PfAMA1, Pb_PfCSP, and Pb_PfCSP+PfAMA1 by dissection at 20 to 22 days post-feeding. Subsequently 5000 sporozoites were injected into 4-week-old C57BL/6 mice (n=5 per group). After 44 h, 100 µl of 5.5 mM AkaLumine-HCl substrate (Fujifilm Wako Pure Chemical Corporation, Osaka, Japan) at a concentration of 5.5 mM was administered intraperitoneally under anesthesia. After 10 minutes, bioluminescence signals were captured using a Newton 7.0 in vivo imaging system (Vilber, Collegien, France). The images were further analyzed using the Kuant program to quantify the bioluminescence signals within mouse livers corresponding to the parasite burden [25]. The average background noise, which was measured from a non-target abdominal region, was eliminated by subtracting it from the average target signal. Sporozoite infectivity was further confirmed by measuring parasitemias in the same mice from days 3 to 5 post-inoculation by flow cytometry using a MACSQuant VYB, as described above.

### Inhibition of PfAMA1-PbRON complex formation by R1 peptide

Enriched fully matured schizonts (10^5^) of chimeric Pb_PfAMA1 were injected intravenously into 5-week-old female BALB/c mice (n=5) along with R1 peptide (VFAEFLPLFSKFGSRMHILK, [31] or control peptide F5 (GDVWLFKTSTSHFAR, [32]) at a dose of 200 μg. R1 peptide or F5 peptide injection was repeated after 1 h, since the stability of R1 peptide diminishes rapidly in the serum. Parasitemias were examined by flow cytometry from 24 h until 96 h post-inoculation, to measure the inhibitory effects of R1 peptide on merozoite infection of erythrocytes. To assess inhibition by R1 peptide during sporozoite infection of hepatocytes, 2000 sporozoites collected from salivary glands were inoculated onto a human hepatoma cell line, HepG2 (100,000 cells/well), together with 1 mg/ml R1 or F5 peptide in 8-well glass Nunc Lab-Tek II chamber slides (Thermo Fisher Scientific). The cells were incubated for 48 h in Dulbecco’s Modified EAGLE Medium with L-glutamine and phenol red (FUJIFILM Wako Pure Chemical) containing 10% FCS, 1% penicillin and streptomycin solution (FUJIFILM Wako Pure Chemical Corporation; 100 IU/mL penicillin, 100 µg/mL streptomycin), 1 µg/mL Amphotericin B (FUJIFILM Wako Pure Chemical Corporation) and 10 µg/mL gentamicin (FUJIFILM Wako Pure Chemical Corporation) at 37°C in the presence of 5% CO_2_. The number of liver stage parasites and their size were evaluated using the tiling images acquired by Nikon Digital Sight 10 camera and Nikon NIS-Elements software (Nikon, Tokyo, Japan).

### Immunization of rats with Pb_PfAMA1 parasites and detection of their inhibitory effects on Pf parasite proliferation in vitro

Four-weeks-old female Wistar rats (n=3 per group) were immunized using a modified protocol based on that described by Friesen et al [47]. Briefly, rats were intravenously injected with 1×10^5^ iRBCs of GFP-Akaluc or Pb_PfAMA1 chimeric parasite line. Once parasitemias reached 0.5%, rats were treated with pyrimethamine in their drinking water at a concentration of 70 µg/ml for 5 days to clear blood stage parasites. Blood samples were collected after the first immunization. Immunization with live parasites was repeated at 12 days after the first inoculation. At 13 days after the second immunization, the rats were terminally exsanguinated by cardiac puncture. Blood samples were allowed to clot for 30 min at room temperature and centrifuged at 2000 x *g* for 10 min at 4°C to collect sera. Sera were sterilized by filtration through a 0.22 μm Millex-GV syringe filter unit (Millipore, Burlington, MA, USA) and heat-inactivated at 56°C for 30 min before use in a growth inhibition assay (GIA). The GIA was performed using human erythrocytes infected with late trophozoites and schizonts stages of *P. falciparum* 3D7 and HB3B strains, one day after synchronization with 5% sorbitol, as described with some modifications [48]. Briefly, 100 µl of synchronous *P. falciparum* parasites were seeded into clear flat-bottom Falcon 96-well tissue culture-treated plates (Corning, NewYork, USA), adjusted to 2% hematocrit and 0.1% parasitemia, and then incubated together with 20 µl of rat serum for 72 h. Parasitemias were evaluated daily by the flow cytometric staining method described above.

### Statistics

Statistical analyses were performed using Graphpad Prism version 9.0.0 (Graphpad, USA). Before statistical analysis, to compare two or more populations, the Kolmogorov-Smirnov test for normality was used to assess whether the values followed a Gaussian distribution. When comparing two groups, the Mann-Whitney test was used for non-parametric data, and the unpaired *t*-test was used for parametric data.

### Ethical statement

The animal study was approved by the Animal Experiment Committee of the Institute of Science, Tokyo. The study was conducted in accordance with the local legislation and institutional requirements.

## Acknowledgments

We are grateful to Dr. Thomas Templeton for critical reading of the manuscript. Rabbit anti-PfAMA1 antiserum was kindly provided by Prof. Takafumi Tsuboi and Prof. Eizo Takashima, Proteo-Science Center, Ehime University. The following reagents were obtained through BEI Resources, NIAID, NIH: Monoclonal Anti-*Plasmodium falciparum* Circumsporozoite Protein (PfCSP), Clone 2A10 (produced in vitro), MRA-183A, contibuted by Elizabeth Nardin, Monoclonal Anti-*Plasmodium* Apical Membrane Antigen 1, Clone 28G2 (produced in vitro), MRA-897A, contributed by Alan W. Thomas; *Plasmodium falciparum*, Strain 3D7, MRA-102, contributed by Daniel J. Carucci; and *Plasmodium falciparum*, strain HB3B, MRA-1227, contributed by Alfred Cortés. Immunoelectron microscopic analysis was supported by the Division of Medical Research Support, the Advanced Research Support Center (ADRES), Ehime University. We also thank Tokiko Okamura for rearing mice and mosquitoes, and technical support.

## Funding

This work was supported by JSPS KAKENHI under Grant number 25K02487 to TI, 24K02272 to NS and 25K18787 to YK; by AMED under Grant Number JP24wm0325074 to TI and JP26wm0325083 to NS; JST SPRING, Grant Number JPMJSP2120 to TS.

## Author contributions

NS, MT and TI: Conceptualization and Study design. QI, NS, YK, TS, MT, HA and TI: Investigation (performed experiments). QI, NS, YK and TI: Data curation. QI, NS, MT and TI: Writing – original draft, and Writing – review & editing.

## Disclosure of conflict of interest

The authors declare no commercial or financial conflict of interest.

## Supplementary Information

**S1 Table. Primers used in this study**

**S1 Fig. Sequence alignment of AMA1 and its interacting region in RON2** (A) Amino acid sequence alignment of AMA1 across *Plasmodium* spp. The protein sequences of AMA1 orthologs from *Plasmodium berghei* (PBANKA_0915000), *Plasmodium falciparum* (PF3D7_1133400), *Plasmodium vivax* (PVP01_0934200), *Plasmodium knowlesi* (PKNH_0931500), and *Plasmodium gallinaceum* (PGAL8A_00360900) were obtained from PlasmoDB (release 68, https://plasmodb.org/) and aligned by Clustal Omega. *, identical amino acid; : and ., amino acid with strong and weak similarity. An arrowhead indicates the proteolysis sites determined in *P. falciparum* AMA1 [27]. Orange and purple arrows indicate domains I and II. Crucial amino acids for RON2 binding are indicated by blue highlights [15]. Green highlighted amino acids were reported to be located at the hydrophobic trough interacting with RON2 [10]. (B) Sequence alignment of RON2 at its C-terminus region between *P. berghei* and *P. falciparum*. Blue highlight indicated the key residues for interaction with PfAMA1 [15].

**S2 Fig. PfAMA1detection at the apical end of Pb_PfAMA1 merozoites by IFA** (A) Acetone-fixed schizonts were incubated with anti-PfAMA1 antiserum (1:2000, shown in red) together with anti-PbRON12 antibodies (1: 1000, shown in green). The merged images also show nuclei stained with Hoechst (shown in blue). The PfAMA1 signal partly overlaps with PbRON12, indicating the close proximity of micronemes to rhoptries. Bar, 5 µm.

**S3 Fig. PfAMA1 complements the roles of PbAMA1 during blood stage parasite proliferation (A)** The reproducibility of parasitemias after BALB/c mice were inoculated with chimeric blood stage parasites. Infected erythrocytes (1 x 10^5^) of parental (GFP-Akaluc)- or Pb_PfAMA1-parasites were inoculated intravenously into five BALB/c mice. The mean parasitemias with standard deviations are plotted, indicating that PfAMA1 functions during *P. berghei* merozoite invasion of erythrocytes.

**S4 Fig. Parasite invasion is inhibited by R1 peptide** (A) R1 peptide decreased merozoite invasion in vivo, confirming the reproducibility of Figure 5. Pb_PfAMA1 schizonts (1 x 10^5^) together with 200 µg R1 or F5 peptide were inoculated intravenously into five BALB/c mice, followed by an additional 200 µg of peptide inoculated 1 h later. Parasitemias at 48 h and 72 h post inoculation are plotted, showing that R1 peptide significantly decreased merozoite invasion (Unpaired t-test, **: P<0.01). (B) LS areas at 48 h after sporozoite and peptide inoculation onto HepG2 cells are plotted, measured by the samples shown in Fig 5B. One hundred-fifty LS selected randomly after R1 or F1 peptide treatment were measured by NIS-Elements software. Parasites develop normally after R1 peptide incubation, as the areas are comparable to those after F5 peptide treatment.

**S5 Fig. Examples of Hoechst/DHE-based dual-color measurement of parasitemias in *P. falciparum* cultures** The dual-color method was performed using blood samples from *P. falciparum* cultures. (A, B) Debris and doublets were excluded using an erythrocyte gate on the forward scatter/side scatter (FSC-A/SSC-A) dot plot (A) and a singlet gate on the FSC-A/FSC-H dot plot (B), respectively. (C) Parasitemias of ring, trophozoite, and schizont stages immediately after sorbitol synchronization (left), at 16 h (middle), and 22 h (right) after synchronization. Each developmental stage was defined by the indicated gates on the ethidium/Hoechst dot plots.

**S6 Fig. Evaluation of the generation of PfAMA1 and PfCSP expressing transgenic parasites** (A) Schematic representation of PfAMA1-replaced (Pb_PfAMA1) and PfCSP-replaced (Pb_PfCSP) locus. Primers used for genotyping PCR are indicated by arrows. (B) PCR genotyping demonstrating that PbAMA1 and PbCSP coding regions were replaced with their orthologous gene expression cassettes. Mouse blood infected with Pb_PfAMA1, Pb_PfCSP, and Pb_PfCSP+PfAMA1 was directly used as templates. Primer sets that specifically amplifying PfAMA1 (upper) and PfCSP (lower) replaced regions were used (see S1 Table). From Pb_PfCSP+PfAMA1-infected mouse blood, both PfCSP- and PfAMA1- related DNA fragments were specifically amplified at expected sizes. (C) IFA demonstrates that PfCSP is expressed on the surface of Pb_PfCSP+PfAMA1 oocyst-derived sporozoites.

## Notes

### Competing Interest Statement

The authors have declared no competing interest.

